# Universality of cell differentiation trajectories revealed by a reconstruction of transcriptional uncertainty landscapes from single-cell transcriptomic data

**DOI:** 10.1101/2020.04.23.056069

**Authors:** Nan Papili Gao, Olivier Gandrillon, András Páldi, Ulysse Herbach, Rudiyanto Gunawan

## Abstract

We employed our previously-described single-cell gene expression analysis CALISTA (Clustering And Lineage Inference in Single-Cell Transcriptional Analysis) to evaluate transcriptional uncertainty at the single-cell level using a stochastic mechanistic model of gene expression. We reconstructed a transcriptional uncertainty landscape during cell differentiation by visualizing single-cell transcriptional uncertainty surface over a two dimensional representation of the single-cell gene expression data. The reconstruction of transcriptional uncertainty landscapes for ten publicly available single-cell gene expression datasets from cell differentiation processes with linear, single or multi-branching cell lineage, reveals universal features in the cell differentiation trajectory that include: (i) a peak in single-cell uncertainty during transition states, and in systems with bifurcating differentiation trajectories, each branching point represents a state of high transcriptional uncertainty; (ii) a positive correlation of transcriptional uncertainty with transcriptional burst size and frequency; (iii) an increase in RNA velocity preceeding the increase in the cell transcriptional uncertainty. Finally, we provided biological interpretations of the universal rise-then-fall profile of the transcriptional uncertainty landscape, including a link with the Waddington’s epigenetic landscape, that is generalizable to every cell differentiation system.

## MAIN

Cell differentiation is the process through which unspecialized stem cells become more specialized. Because of its importance in development, cellular repair, and organismal homeostasis, the molecular mechanisms of cell differentiation has been the subject of intense scrutiny. Since roughly 50 years ago – along with the promulgation of the central dogma of molecular biology by Francis Crick and the characterization of the lactose operon by François Jacob and Jacques Monod – the existence of a genetic program has become a prevailing explanation for the cell differentiation process. Although the details were originally not defined, at least not formally, such a genetic program purports a constellation of master genes (i.e., transcription factors) that orchestrate the transcription of downstream target genes in a precise spatiotemporal fashion, resulting in long-lasting alterations in the gene expression patterns ^1–3^. A notable experimental evidence substantiating this view is the overexpression of myoD inducing a myogenic phenotype in seemingly naive cells ^4^. Over the past few decades, the repertoire of such master genes across numerous stem cell systems, such as Nanog, Oct4, Sox2, BATF and MyoD, begin to coalesce ^5–7^.

Recent advances in single-cell technologies has revealed new aspects of the cell differentiation that are incompatible with the idea of ordered and programmed (i.e., deterministic) gene expression. More specifically, single-cell data paint a stochastic differentiation process that increases cell-to-cell variability of gene expression. Such an observation has been made for a wide variety of cell differentiation systems, including chicken erythroid progenitors ^8^, erythroid myeloid lymphoid (EML) cells ^9^, mouse embryonic stem cells (mESCs) ^10,11^, and human CD34+ cells ^12^. Interestingly, a similar increase of gene expression variation was also observed during the de-differentiation of somatic cells into iPSCs ^13^. Stochastic gene expression also appears to have functional role beyond cell differentiation systems. For example, an increase in cell-to-cell variability of gene expression has been reported during a forced adaptation of budding yeast cells to unforeseen challenges ^14^.

Based on the observations of single-cell data above, a different view of cell differentiation begins to materialize. Instead of stem cells following an identical genetic program, the cell differentiation is akin to a dynamic exploratory process. More specifically, in this view, the cell differentiation is thought to proceed as follows ^14–17^:

I. extrinsic and/intrinsic internal stimuli, such as a medium change or the addition of new molecules in the external medium, trigger a cellular response that destabilizes the initial high potency cell state;
II. each cell alters its internal cell state and engages an exploratory dynamics through a combination of the inherent stochastic dynamics of gene transcription and the emergence of new stable cell state(s). At the cell population level, we observe a rise in the cell-to-cell variability of gene expression;
III. a physiological selection / commitment to one stable lineage among possibly multiple lineages that arise from the degeneracy of the system;
IV. finally, a reduction in the exploratory dynamics commences along with the establishment of stable cell state(s) corresponding to differentiated cell type(s).

The above view is compatible with the idea that cell phenotype transition results from the dynamics of an underlying stochastic molecular network ^18,19^. In 1957, Conrad Waddington proposed the presently well-known epigenetic landscape that likens the cell differentiation process to a ball rolling on a downward sloping surface, starting from a state of high cell potency and ending at one of possibly several states of low cell potency. The landscape itself is shaped by the action of the genes and gene network – depicted in the less-frequently-shown part B of Waddington’s original figure as a network of ropes that are tied to the surface, creating valleys and hills. Although the epigenetic landscape was originally proposed only as a metaphor of how gene regulation governs the cell differentiation process, this landscape has been formalized within the framework of dynamical systems theory ^20^. The valleys in the Waddington’s epigenetic landscape are equated to stable states of a dynamical system, called attractors, while the hills are often interpreted as energetic barriers.

The Waddington’s metaphor has further been re-examined in a series of works, where the behaviour of the underlying dynamical gene regulatory network (GRN) drives cell fate determination. Some of these studies employed stochastic simulations of simple gene networks with bi-/tri-stable states to describe the role of non-genetic cell heterogeneity in cell differentiation processes ^21–23^. Here, to quantitatively reconstruct the epigenetic landscape, an underlying GRN structure must be assumed *a priori*. In reality, the GRN driving cell fate determination is complex and proved to be challenging to infer from data ^6,24–28^. Even when the complete regulatory system is known, the high-dimensionality of the parameter space makes the landscape generation computationally prohibitive ^21^, especially in the absence of an analytical solution, which are slowly emerging ^29^. Thus, the above approaches provide more of a conceptual understanding than a mechanistic or molecular explanation.

A number of recent studies provided a graphical representation of the differentiation process based on single-cell transcriptomic data that conforms with the Waddington’s epigenetic landscape ^30–34^. More specifically, these studies reconstructed the epigenetic landscape from single-cell gene expression data using probabilistic and quasi-potential methods, for example by applying Hopfield neural networks ^30,31^, a cell-density based strategy ^33^, network entropy measurements 34 or more recently Large Deviation Theory ^29^. However, with the exception of Fard et al. ^31^ and Lv et al., 2014 ^29^, the aforementioned studies produced monotonic descent passages during cell differentiation, mimicking closely the Waddington’s epigenetic landscape metaphor (see for example ^22,34^). Also, none of the above studies consider directly the cellular mechanism that generates stochastic gene transcriptional bursts.

In the present work, we aimed to shed light on the gene transcriptional mechanism behind the rise-then-fall trajectory of cell-to-cell variability in gene expression observed during the cellular differentiation process. To this end, we analyzed a collection (8) of published single-cell transcriptomic datasets from various cell differentiation systems, comprising both single-cell RT-qPCR (scRT-qPCR) ^8,10,12,35–37^ and single-cell RNA-sequencing (scRNA-seq) ^38,39^. We employed a likelihood-based analysis using a recent method CALISTA (Clustering And Lineage Inference in Single-cell Transcriptomics Analysis) ^40^. The analysis relied on a mechanistic model of the stochastic gene transcriptional bursts to characterize single-cell gene expression distribution. We defined a new concept, called transcriptional uncertainty landscape, based on the cell likelihood value from CALISTA analysis to characterize the stochastic dynamics of the gene transcription process during cell differentiation. The stochastic gene transcriptional model enabled identifying the specific parameters or mechanisms that explain the observed changes in the the gene transcriptional uncertainty at the single-cell level ^41^. For two additional single-cell datasets, we also evaluated the single-cell RNA-velocity using the recently published Velocyto method^42^. The two-state model parameter analysis, combined with RNA-velocities, provided insights into the mechanism regulating cell fate decisions, specifically on the role of stochastic gene transcriptions in the differentiation processes and on the possible mechanism generating this stochasticity.

## RESULTS

### Single-cell Transcriptional Uncertainty Landscape using CALISTA

In this work, we used CALISTA, a likelihood-based bioinformatics toolbox designed for an end-to-end analysis of single-cell gene expression data, to evaluate the transcriptional uncertainty of each individual cell based on its gene expression data ^40^. CALISTA uses the two-state model of stochastic gene transcription bursts to characterize the distribution of mRNA counts in individual cells ^43^. In the model, a gene promoter stochastically switches between ON and OFF state, and only in the ON state can gene transcription occur. The distribution of mRNA depends on four model parameters: *θ*_*on*_ (the rate of promoter activation), *θ*_*off*_ (the rate of promoter inactivation), *θ*_*t*_ (the rate of mRNA production when the promoter is on the ON state), and *θ*_*d*_ (the rate constant of mRNA degradation) ^27,44^(see Fig. 1). For example, when *θ*_*off*_ ≫ *θ*_*on*_ and *θ*_*off*_ ≫*θ*_*d*_, keeping *θ*_*t*_/*θ*_*off*_ fixed, mRNA are produced through bursts of short but intense transcription, which is a typical case observed for gene transcriptions in single cells ^45,46^.

**Figure 1.**
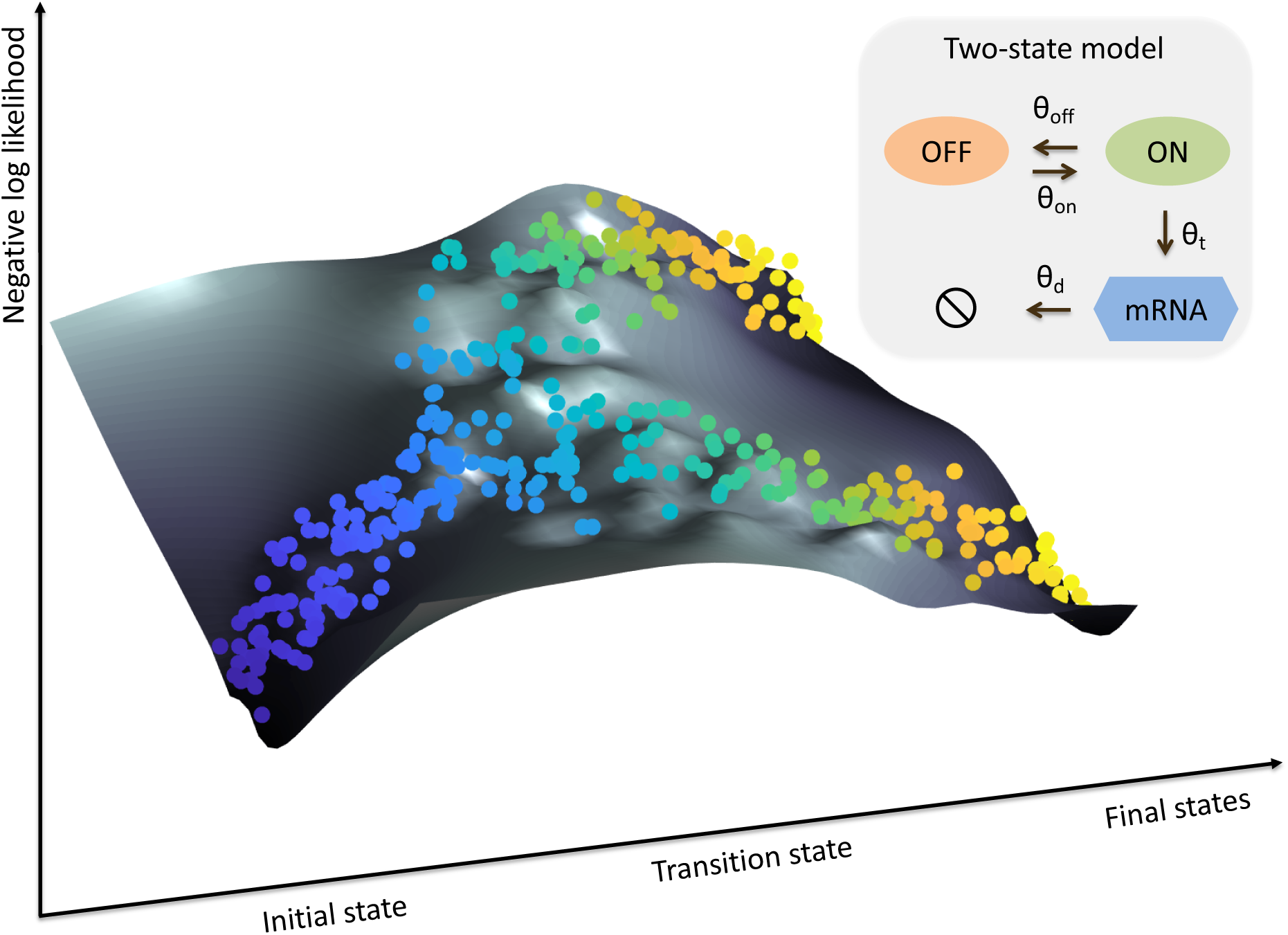
Single-cell transcriptional uncertainty landscape. The illustration depicts the landscape of single-cell transcriptional uncertainty during a differentiation process over the (pseudo) time (from blue to yellow). Each dot corresponds to a cell in the single-cell transcriptomic dataset. Cells start their journey from a valley in the landscape, through a hill, before ending at one of the final valleys / states.

Our analysis procedure using CALISTA involves four main steps: cell clustering, lineage inference, and calculation of single-cell transcriptional uncertainty. In the analysis, each cell is assigned a likelihood value, which is computed from the probability of its gene expression (mRNA counts) based on the mRNA distribution given by the two-state model above. In the single-cell clustering, we employed a greedy algorithm to find single-cell clusters that maximizes the total likelihood value for the cells. In the lineage progression inference, we connected cell clusters sharing similar gene expression distributions – by computing cluster distances based on cell likelihood – to form a hierarchical network. Here, we assigned each cell to an edge in the lineage progression network that is pointing to or emanating from its respective cluster, such that its single-cell likelihood is maximized. Finally, we computed single-cell transcriptional uncertainty as the negative logarithm of the single-cell likelihood (NLL) value. The full detail of CALISTA can be found in a recent publication ^40^.

The single-cell likelihood value reflects the joint probability of its gene expression repertoire. A cell with a low likelihood value may indicate that the gene expression of the cell is different from its neighboring cells, i.e. the cell is an outlier. But, more interestingly, a low likelihood value may also correspond to a cell state of high uncertainty in the gene expression. The group of cells in such high uncertainty state have gene expressions that are dissimilar to each other, and thus, the gene expression distributions will have high entropy. As mentioned above, in this work we used the negative logarithm of the single-cell likelihood value as a metric of single-cell transcriptional uncertainty. By plotting the single-cell transcriptional uncertainty over the two-dimensional projection of the single-cell transcriptomics data – for example, using the first two principal components from Principal Component Analysis (PCA), we constructed a transcriptional uncertainty landscape in the form of a surface plot of the NLL values. By visualizing the single-cell transcriptional uncertainty as a surface plot, we can study the landscape of transcriptional uncertainty during cell differentiation at single-cell resolution. On the single-cell transcriptional uncertainty surface, an aberrant cell can be easily distinguished from a cell of high uncertainty state, since an aberrant cell will appear isolated from its nearby cells and will be located on a region with high NLL.

### Transcriptional uncertainty landscape of iPSC cell differentiation to cardiomyocytes

In the following, we demonstrated an application of our procedure described above to a single-cell transcriptional dataset from cardiomyocytes differentiation from human induced pluripotent stem cells (iPSCs) ^35^. The single-cell clustering of CALISTA returned five clusters ^40^, in good agreement with the number of cell types reported in the original study. CALISTA identified one bifurcation event in the lineage progression, which led to two cell lineages ^35^. Fig. 2a-b gives the single-cell transcriptional uncertainty landscape showing cells exiting the initial epiblast state that is characterized by a valley in the landscape, passing through a hill of high transcriptional uncertainty corresponding to primitive streak (PS)-like progenitor state, before ending up at one of the low transcriptional uncertainty terminal states corresponding to either mesodermal (desired) or endodermal (undesired) fate. As depicted in Fig, 2c, the intermediate cell cluster (cluster 2) comprising PS-like cells have higher cell uncertainty (lower single-cell likelihood) than the other clusters. Figures 2d and 2e give the moving-averaged uncertainty values for pseudotemporally ordered cells using a moving window of 10% of the total cells for both endodermal and mesodermal paths, respectively. The moving-averaged transcriptional uncertainty for the two differentiation paths follows a rise-then-fall trajectory where the peak of uncertainty coincides with the lineage bifurcation event.

**Figure 2.**
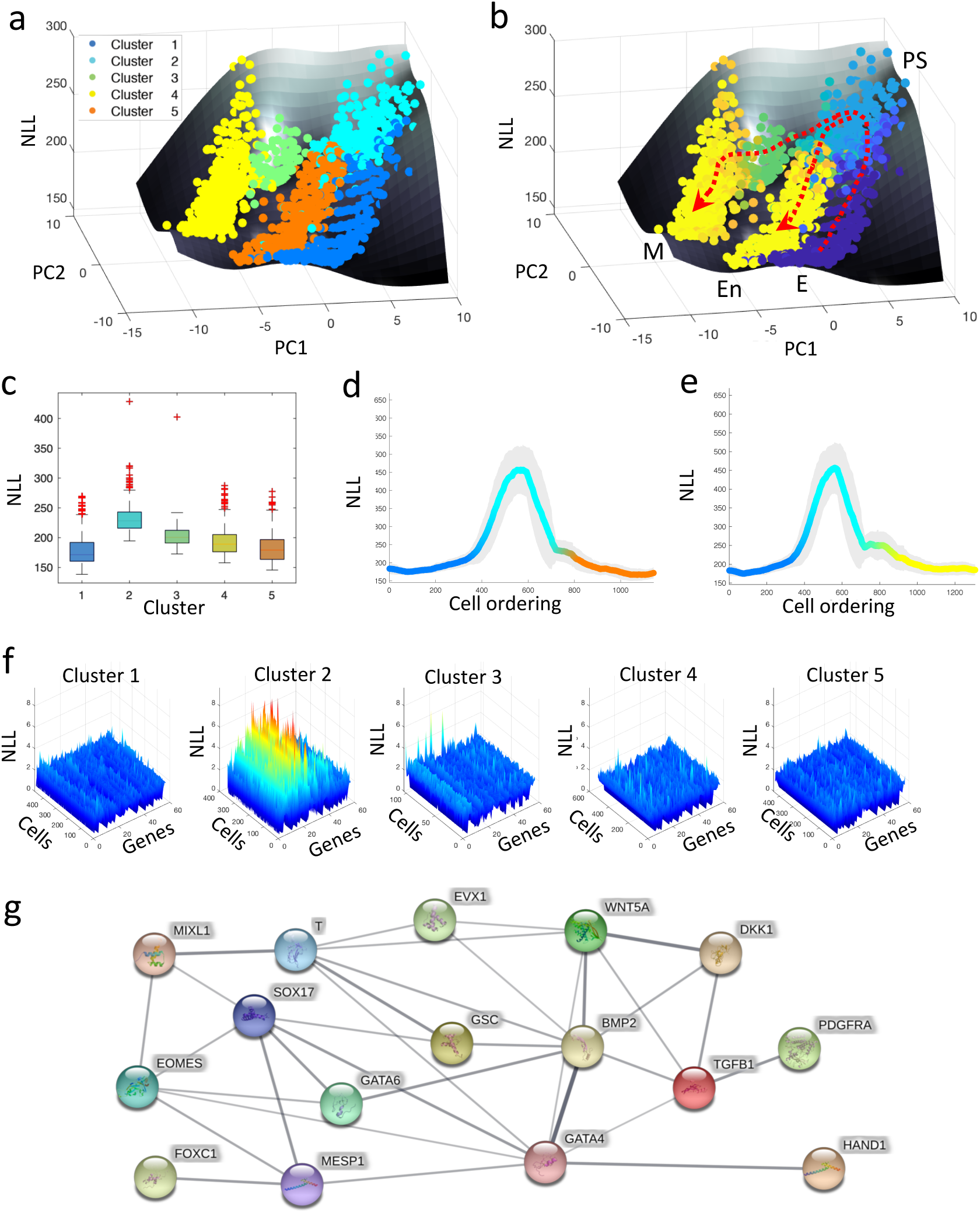
Analysis of single-cell transcriptional profiles during iPSC differentiation into cardiomyocytes. The single-cell gene expression dataset was taken from the study of Bargajeet al. ^35^ (a-b) Epigenetic landscape plots estimates by CALISTA. Each dot on the landscape represents a cell where the colors indicate (a) the cell cluster, (b) pseudotime (from dark blue to yellow). The x-y axes of the landscape plots correspond to the first and second principal component (PC) coordinates, respectively, of the single-cell transcriptomic data. (c) Boxplots of NLL values for each single-cell cluster. (d-e) Moving-window average NLL along (d) endoderm and (e) mesoderm fate trajectory. (f) NLL of each gene and cell for every single-cell cluster. (g) Protein-protein interaction network of top variable genes inferred by STRING ^49^. Blue nodes represent transcription factors, while red nodes denote proteins involved in signal transduction. The width of the edges denotes the confidence for the inferred relationship (thicker edge = higher confidence).

We explored whether the rise-then-fall in uncertainty is an artefact from using the two-state model to evaluate the cell likelihood values. To this end, we implemented a modified version of the algorithm for ordering cells by calculating the cell likelihood values using the empirical (observed) distribution, instead of the analytical distribution from the two-state model. As shown in Supplementary Fig. S1, the transcriptional uncertainty landscape from the modified implementation shows a strong resemblance to the original one. We also investigated whether the number of clusters may affect the landscape, in which using too few of clusters may artificially inflate the uncertainty due to mixing of cells from different states. We reran CALISTA by using a higher number of clusters (set to nine based on the eigengap heuristic ^47^). The hill in the uncertainty landscape is again seen around the bifurcation event upon using a higher number of cell clusters (Supplementary Fig. S2). Finally, we used a different algorithm to cluster cells, specifically using a Laplacian-based clustering algorithm SIMLR ^48^, to test whether the shape of the transcriptional uncertainty landscape changes with the clustering algorithm. The single-cell clusters can be interpreted as the transitional states that the differentiating cells go through. Starting with the result of SIMLR cell clustering, we then generated the lineage progression and estimated the cell likelihood values using CALISTA. The transcriptional uncertainty landscape from SIML cell clustering has the same shape as that in Fig. 2a-b, demonstrating that the transcriptional uncertainty landscape observed above is not dependent on using CALISTA for cell clustering (Supplementary Fig. S3).

To further elucidate the role of specific genes in shaping the transcriptional uncertainty landscape, we looked at the transcriptional uncertainty associated with individual genes. Fig. 2f depicts the NLL distribution of each gene for the five single-cell clusters. As expected, cell in cluster 2 have generally higher NLL than those in the other clusters. Fig. 2f clearly illustrates that within cluster 2, some genes show higher NLL values than the others (Supplementary Figure S4). To identify the important genes related to transcriptional uncertainty, we identified genes with NLL values exceeding a threshold δ for at least 30% of the cells in each cluster, where *δ* is set to 3 standard deviation above the overall mean NLL for all cells and genes in the dataset (see Methods Eq. (2)). None of the genes in clusters 1, 4 and 5 have NLL above the threshold. Meanwhile, 16 and 8 genes in clusters 2 and 3, respectively, pass the above criterion for high uncertainty with 4 common genes between the two gene sets (Supplementary Table S1). Genes with high transcriptional uncertainty in cluster 2 may have functional roles in cell fate determination. The gene set of cluster 2 includes known genes upregulated only in the PS-like state (e.g. EOMES, GSC, MESP1 and MIXL1), as well as markers of mesodermal and endodermal cells (e.g. BMP4, HAND1, and SOX17) ^35,40^ (Supplementary Figure S5). Meanwhile, the main contributors to cell uncertainty in cluster 3 (e.g. BMP4 and MYL4 ^35,40^) are known transition genes between PS-like cells and the final mesoderm fate (Supplementary Figure S6). Fig. 2g depicts the protein-protein interaction (PPI) network related to the gene set of cluster 2 using STRING (minimum required interaction score of 0.4) ^49^, indicating that these genes form a strongly interconnected hub of known transcription factors and molecules involved in the signal transduction of embryonic development (Supplementary Table S1).

### Transcriptional uncertainty landscapes of cell differentiation

We further applied the procedure above to seven additional single-cell transcriptomic datasets that were generated using scRT-qPCR ^8,10,12,36,37^ and scRNA-sequencing ^38,39^, to assess the universality of the rise-then-fall feature of single-cell transcriptional uncertainty landscape during cell differentiation. The first of these datasets came from 405 cells during mouse embryonic fibroblast (MEF) reprogramming into induced neural (iN) and myogenic (M) cells ^38^. Like the iPSC differentiation above, the lineage progression has a single bifurcation point. As depicted in Figure 3a, the single-cell transcriptional uncertainty increases from the initial MEF state and reaches a peak around the bifurcation before decreasing toward two end-point cell fates. The rise-then-fall of transcriptional uncertainty in the MEF reprogramming is in good agreement with what we observed in the iPSCs differentiation above. Higher entropy of gene expression distribution in a cell population has also been reported in the reprogramming of iPSCs^13^.

**Figure 3.**
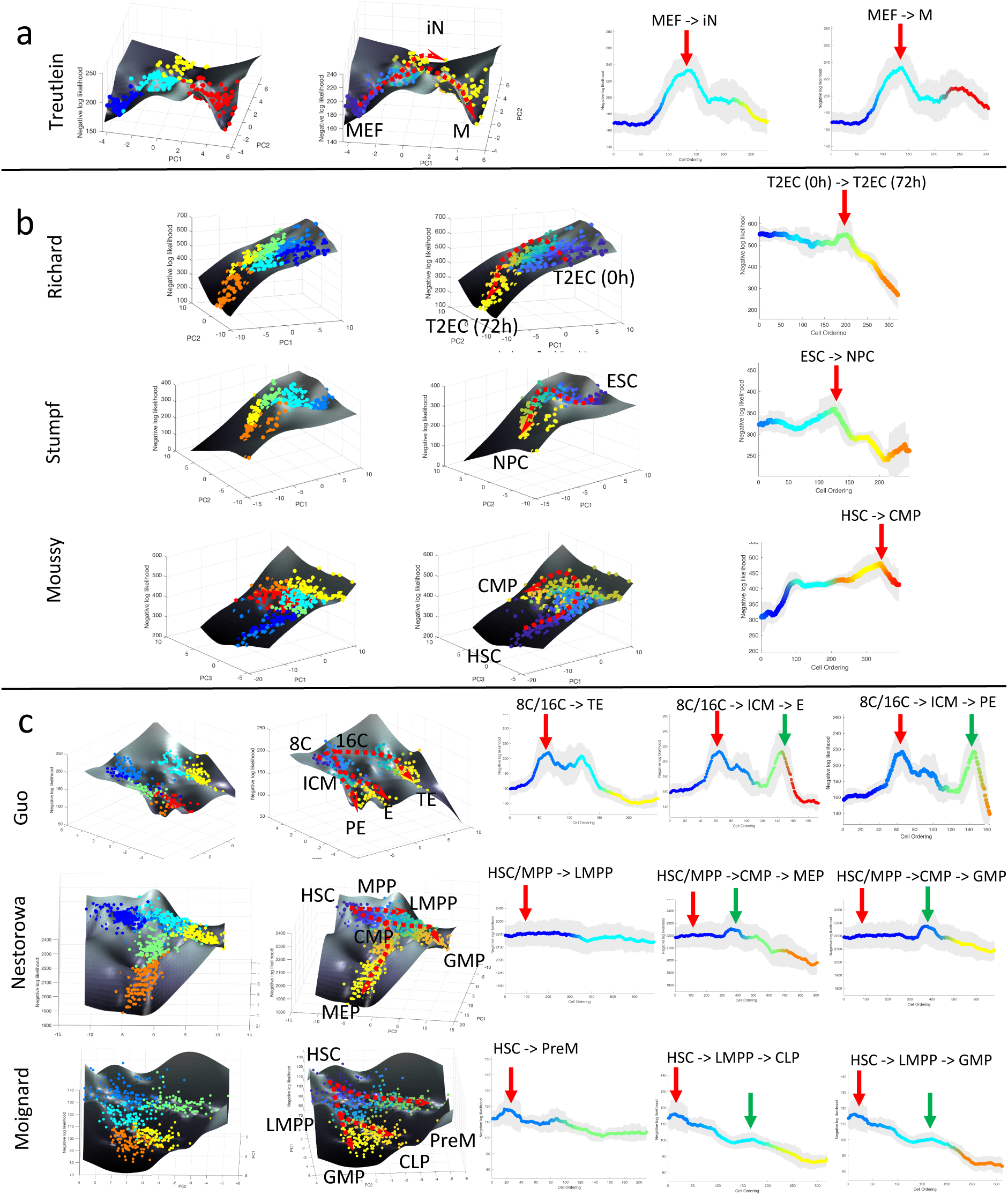
CALISTA analysis of single-cell expression data. (a-c) Landscape plots (based on cell clusters and pseudotime) and moving-averaged NLLs for each differentiation path of (a) single-branching trajectory (Treutlein dataset ^38^), (b) linear trajectories (Richard ^8^, Stumpf ^10^, and Moussy ^12^ datasets), (c) multi-braching trajectories (Guo ^36^, Nestorowa ^39^, and Moignard ^37^ datasets). Green and red vertical arrows in moving-averaged NLL plots indicate the first and second peak in cell uncertainty, respectively.

Next, we analyzed datasets from cell differentiation processes without a lineage bifurcation and with multiple lineage bifurcations. Three scRT-qPCR datasets came from differentiation systems without bifurcation, including Richard et al. study on chicken erythrocytic differentiation of T2EC cells ^8^, Stumpf et al. study on differentiation of mouse embryonic stem cells (ESC) to neural progenitor cells (NPC) ^10^, and Moussy et al. study during CD34+ cell differentiation ^12^. The single-cell clustering and lineage progression by CALISTA produced the expected cell differentiation trajectory (see Supplementary Figure S7-9). The single-cell transcriptional uncertainty landscapes of these three differentiation systems, as shown in Figure 3b, exhibit a rise-then-fall profile, creating a hill that the cells traverse through in the differentiation process. A transitory increase in single-cell gene expression uncertainty was reported either directly or indirectly in the original publications. In Richard et al. ^8^ and Stumpf et al ^10^, the authors adopted the Shannon entropy to quantify cell-to-cell variability (uncertainty), while Moussy et al. ^12^ reported an unstable transition state with ‘hesitant cells’ flipping their morphology between polarized and round shapes before committing to the common myeloid progenitors-like fate. Morphological uncertainty therefore corresponded to a higher transcriptional uncertainty. Note that Moussy et al. study looked at only the initial phase of the (hematopoietic) cell differentiation, and thus, it is likely that the differentiation process had not completed for the cells in the dataset.

The next set of single-cell gene expression data came from differentiation systems with multi-branching lineage, including Guo et al. study during mouse embryo development from zygote to blastocyst ^36^, Nestorowa et al. ^39^ and Moignard et al. ^37^ studies on hematopoietic stem cell differentiation. Figure 3c show the single-cell transcriptional landscape for each of the datasets. For Guo et al. study, we identified 7 cell clusters and identified two bifurcations in the lineage. Here, we observed two hills in the transcriptional uncertainty landscape, each coinciding with a bifurcation event in the lineage progression – one at 32-cell stage (cluster 2 to cluster 3 and 4) and another at 64-cell stage (cluster 4 to cluster 6 and 7) (see Supplementary Figure S10). For Nestorowa et al. ^39^ (Supplementary Figure S11) and Moignard et al. ^37^ (see Methods and ^40^) datasets, we again observed peaks in the transcriptional uncertainty landscape that colocalize with the bifurcation points in the lineage progression.

The use of the two-state mechanistic gene transcriptional model within CALISTA enabled us to probe into a mechanistic explanation for the observed shape of the transcriptional uncertainty landscape. Table 1 show the pairwise Pearson correlations between the cell-averaged NLL of each cluster with two biologically interpretable model parameters, namely transcriptional burst size (number of transcripts generated in each burst) and burst frequency (occurrence of burst per unit time) ^50^ (see Methods). The Pearson correlations indicate that the single-cell gene expression uncertainty increases with higher burst size and burst frequency (*p*-value ≤ 0.01). Higher transcriptional burst size and frequency are associated with a lower *θ*_*off*_ – a lower rate of promoter turning off – and a greater *θ*_*on*_ – higher rate of promoter turning on. One possible explanation for such change in model parameters is a higher chromatin accessibility during the transition period of cell differentiation. This finding is consistent with the view that stem cells increase its gene expression uncertainty or stochasticity by adopting a more open chromatin state to enable the exploration of the gene expression space ^33,50–52^.

**Table 1.**
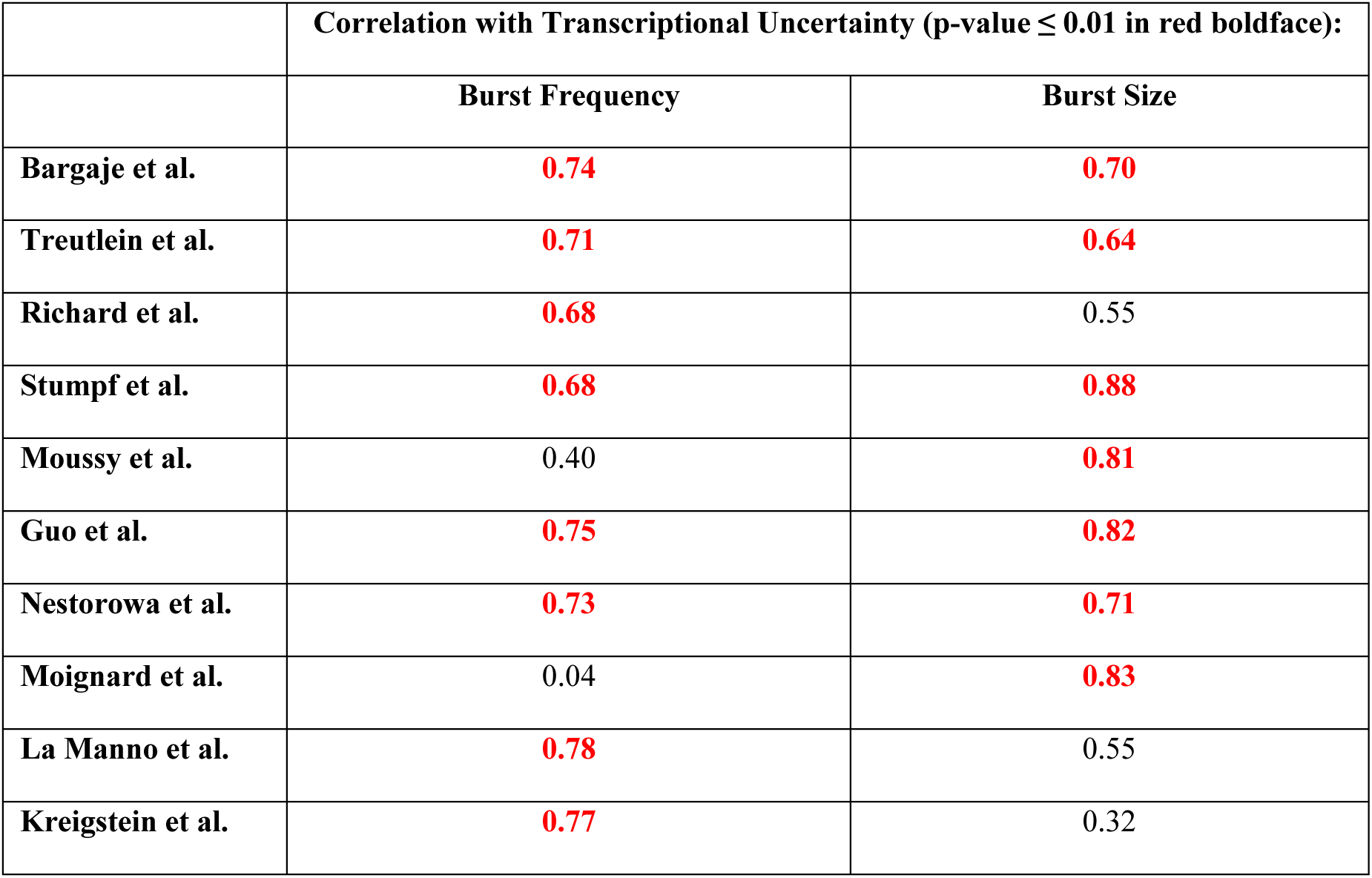
Pairwise correlation coefficients between transcriptional uncertainty and transcriptional burst frequency / burst size.

|  | **Correlation with Transcriptional Uncertainty (p-value $\leq 0.01$ in red boldface):** |  |
| --- | --- | --- |
|  | **Burst Frequency** | **Burst Size** |
| **Bargaje et al.** | **0.74** | **0.70** |
| **Treutlein et al.** | **0.71** | **0.64** |
| **Richard et al.** | **0.68** | 0.55 |
| **Stumpf et al.** | **0.68** | **0.88** |
| **Moussy et al.** | 0.40 | **0.81** |
| **Guo et al.** | **0.75** | **0.82** |
| **Nestorowa et al.** | **0.73** | **0.71** |
| **Moignard et al.** | 0.04 | **0.83** |
| **La Manno et al.** | **0.78** | 0.55 |
| **Kreigstein et al.** | **0.77** | 0.32 |

### Coupling cell uncertainty with RNA velocity

In a recent paper ^42^, La Manno and colleagues introduced the concept of RNA velocity, which involves computing the rate of change of mRNA from the ratio of unspliced to spliced mRNA. A positive RNA velocity indicates an induction of gene expression, while a negative RNA velocity indicates a repression of gene expression. La Manno et al. demonstrated that RNA velocities are able to predict the trajectory of cells undergoing a dynamical transition, such as in circadian rhythms or cell differentiation. In the following, we explored the relationship between RNA velocities and single-cell transcriptional uncertainty.

We evaluated the single-cell transcriptional uncertainty and RNA velocity for two single-cell gene expression datasets that were previously analyzed in La Manno et al. ^42; 42^. The first dataset came from human glutamatergic neurogenesis which has a linear (non-bifurcating) lineage progression. Figure 4 (top row) depicts the cell clustering, single-cell transcriptional uncertainty, and RNA velocities (see also Supplementary Figure S12). The single-cell transcriptional uncertainty landscape again has the rise-then-fall shape, as in the other cell differentiation systems discussed above. Interestingly, the same rise-then-fall profile is also seen in the RNA velocity. As illustrated in Figure 4, the increase and decrease of the RNA velocity preceed the transcriptional uncertainty, and the peak of RNA velocity occurs prior to those of the transcriptional uncertainty (see Supplementary File S1 for an animated illustration). Furthermore, a gene-wise cross-correlation analysis confirms a positive correlation between RNA velocity and single-cell transcriptional uncertainty with a delay for individual genes (see Supplementary Figure S13).

**Figure 4.**
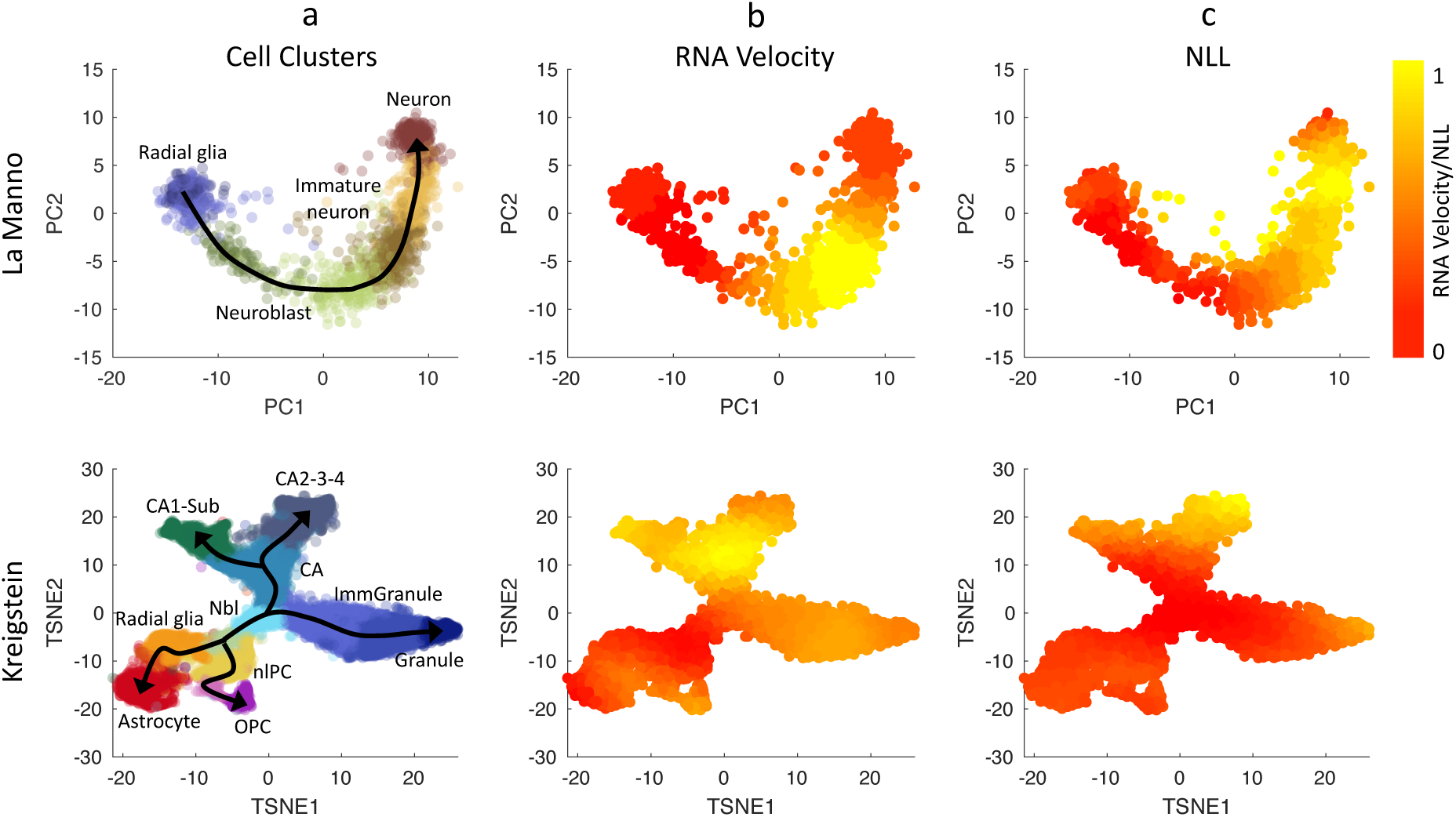
Comparison between RNA Velocities estimated using Velocyto ^42^ and CALISTA NLL values. (Top row) Human glutamatergic neurogenesis in La Manno et al. study ^42^. (Bottom row) Mouse hippocampal neurogenesis in Kreigstein et al. study ^53,53^. (First column) Cell clustering assignments evaluated from Velocyto. Normalized values for Euclidean norm of RNA velocities (2^nd^ column), CALISTA single-cell transcriptional uncertainty (NLL; 3^rd^ column). The colors in the first column indicate the cell clusters, and those in the second-third columns indicate the normalized cell-wise RNA velocities and NLL values respectively.

We also compared RNA velocity and single-cell transcriptional uncertainty for another dataset from mouse hippocampal neurogenesis with a multi-branching lineage ^53^. Figure 4 (bottom row) shows that like in the neurogenesis dataset earlier, the RNA velocity increases and then decreases during cell differentiation, and the change in the RNA precede that of the transcriptional uncertainty (see Supplementary File S2). Also, the RNA velocity peaks take place before the transcriptional uncertainty peaks. The rise-then-fall dynamic of the RNA velocity seen in the two datasets above is consistent with the view that cells engage in an exploratory stochastic dynamics as they leave the progenitor state, and disengage this explorative mode as they reach toward the final cell state.

## DISCUSSION

Although Waddington’s epigenetic landscape was originally proposed only as a metaphor, the landscape has helped stem cell researchers to conceptualize the cell differentiation processes through canalization of cell lineages. As mentioned earlier, much of the existing literature on the analytical reconstruction of the epigenetic landscape relied on either a dynamical system theory applied to a simple gene network, or a thermodynamic interpretation based on the potential energy of a reaction ^21–23^. In our study, we did not make any prior assumptions on the gene regulatory network driving the differentiation process nor on the characteristics of the landscape, such as the existence of a stable valley or that of an energetic barrier (hill). Here, we assumed that the gene transcription at the single-cell level occurs as stochastic transcriptional bursts, in which mRNA counts follow the equilibrium distribution that is given by the two-state gene transcription model ^43^. Therefore, the transcriptional uncertainty landscape in our study is a reflection of the dynamic evolution of gene transcriptional stochasticity within individual cells during the cell differentiation process.

The reconstruction of the transcriptional uncertainty landscapes from 10 single-cell transcriptomic datasets of various cell differentiation processes in our study reveals a universal rise-then-fall trajectory in which cells start from a high potency state with a uniform gene expression pattern in the cell population, then progress through transitional cell state(s) marked by increased transcriptional uncertainty (i.e., higher cell-to-cell variability), and eventually reach one of possibly several final cell states with again a uniform gene expression pattern among the cells. Furthermore, the peaks of the transcriptional uncertainty landscape colocalize with forks in the cell lineage. The rise-then-fall in cell uncertainty agrees well with other reports from different cell differentiation systems ^8–12,54,55^, suggesting that stem cells go through a transition state of high gene expression uncertainty before committing to a particular cell fate. The existence of a hill or barrier during the intermediate stage of cell differentiation has also been proposed in previous studies ^14,31,56^. In particular Moris and colleagues compared this transition state to the activation energy barrier in chemical reactions ^56^. We noted however, that a hill in our transcriptional uncertainty landscape is a reflection of a peak in the cell-to-cell gene expression variability, and thus does not represent a resistance or barrier that a cell has to overcome.

In the analysis of iPSCs differentiation into cardiomyocytes ^35^, the key genes regulating cardiomyocyte differentiation are among the largest contributors to the overall transcriptional uncertainty at or around the peak in the landscape, supporting the idea that dynamic cell-to-cell variability has a functional role in cell-fate decision making processes ^21,57,58^. Such an idea would be in congruence with the recent demonstration that, in a physiologically relevant cellular system, gene expression variability is functionally linked to differentiation ^57,58^.

The rise-then-fall trajectory in the transcriptional uncertainty landscape are more pronounced in some datatsets than in others. For example, in Nestorowa ^39^ and Moignard ^37^ datasets (see Figure 3c), peaks in the transcriptional uncertainty landscape are less noticeable than in the other differentiation systems. We noted that cells in the Nestorowa ^39^ and Moignard ^37^ studies were pre-sorted by using flow cytometry based on the expression of surface protein markers. We posited that at least some cells in the transition state(s) might have been lost during the cell pre-sorting since such cells might not express the chosen surface markers strongly.

Further, the correlation analysis between the cell transcriptional uncertainty and biologically meaningful rates of the stochastic gene transcription model showed strong positive correlations with transcriptional burst size and frequency. In agreement with the result of the correlation analysis, several studies have reported an increase in gene transcriptional bursts during transition states in cell differentiation and other recent studies have suggested that both burst frequency and burst size regulate gene expression levels ^33,51,52^. Importantly, our comparison of the single-cell transcriptional uncertainty and the single-cell RNA velocity revealed that an increase (decrease) in RNA velocity predicts an increase (decrease) in transcriptional uncertainty after a short delay, and that a peak of RNA velocity preceeds that of the transcriptional uncertainty.

The aforementioned observations, while correlative in nature, points to possible biological mechanisms underlying the universal dynamic feature of single-cell transcriptional uncertainty during cell differentiation. At the start of the differentiation process, cells engage an exploratory search dynamics in the gene expression space by increasing stochastic transcriptional burst size and burst frequency. The putative objective of such a stochastic search is to optimize the cell’s gene expression pattern given its new environment. The engagement of this stochastic exploratory mode is supported by the observed increased in the overall RNA velocity and its expected-but-delayed effect in elevating the cell-to-cell gene expression variability (i.e. higher transcriptional uncertainty). Increased transcriptional burst size and frequency suggest a promoter state of stochastic gene transcription that favors the ON state (higher *θ*_*on*_ and lower *θ*_*off*_).

A possible mechanism behind this exploratory search dynamics is an increase in chromatin mobility, driven by metabolic alterations in early differentiation ^59^. Multiple studies have demonstrated that a mismatch between the intracellular state of stem cells and their immediate environment can lead to metabolic reorganization ^60–62^. More specifically, a change in the balance between glycolysis and OXPHOS metabolism has been associated to numerous differentiation processes (see Richard et al. ^63^ and references therein).

Furthermore, changes in the metabolic flux state in early differentiation can modulate the activity of chromatin modifying enzymes through their metabolic co-factors ^64^, or in more direct fashion ^65^ and alter the cell differentiation outcome. A more dynamic state of the chromatin is associated with more variable gene expressions due to the changes in the opening-closing dynamics (breathing) of the chromatin ^66^. As the cells approach the final state, cells disengage the exploratory search mode, as the cells approach an optimal gene expression and metabolic state associated with a chosen cell type.

The findings of our analysis fit within the paradigm of a stochastic stem cell differentiation process as proposed in the introduction. The disordered gene expression pattern during the transition period can be seen as an exploratory dynamics to find the optimal pattern(s) ^14,17^. The transcriptional uncertainty in our analysis can be interpreted as the width of the valley in Waddington’s epigenetic landscape. If one considers the epigenetic landscape as a depiction of the accessible gene expression subspace through with stochastic single-cell trajectories pass during differentiation, a wider valley indicates a more variable gene expression pattern. While in the original Waddington’s epigenetic landscape the valley naturally widens around the branching point in the cell lineage, our analysis shows that a widening of the valley (an increase in transcriptional uncertainty) also occurs in non-branching lineage. In other words, the increase in transcriptional uncertainty appears to be a universal feature of the cell differentiation process, one that arises from the engagement of exploratory mode through increased stochasticity in transcriptional bursts, as explained above.

## METHODS

### Main steps of CALISTA workflow

Herein, we briefly describe the main steps involved in the calculation of single-cell transcriptional uncertainty using CALISTA (for further details see ^40^).

#### Pre-processing

Given an *N* × *G* single-cell expression matrix *M*, where *N* denotes the number of cells and *G* the number of genes, the pre-processing in CALISTA involves two steps: a normalization of the expression data *m*_*n,g*_ – i.e. the number of transcripts of gene *g* in the *n*-th cell, and a selection of the most variable genes ^40^.

#### Cell clustering

CALISTA clustering follows a two-step procedure. The first step involves a greedy optimization strategy to find cell clustering that maximizes the total cell likelihood, i.e. the sum of the likelihood value for all cells. The single-cell likelihood value is computed as the joint probability of the cell’s gene expression data, which is set equal to the product of the probabilities of the mRNA counts for the selected genes based on the mRNA distribution from the two-state stochastic gene transcription model. To avoid issues with numerical overflow, we use the logarithm of the cell likelihood. By performing the greedy optimization multiple times, a consensus matrix containing the number of times two cells in the dataset are put in the same cluster, is generated.. In the second and final step, CALISTA generates the cell cluster assignments by using *k*-medoids clustering based on the consensus matrix. The final outcome of CALISTA’s clustering is the assignment of cells into *K* clusters and the optimal model parameters for the two-state gene transcription model: 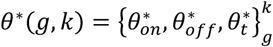 for each gene *g* in cluster *k* (for simplicity, we set 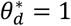 and scale the other parameters by the degradation rate) ^40^.

#### Lineage progression inference

In CALISTA, cell lineage progression is inferred based on cluster distances – a measure of dissimilarity between two clusters. The cluster distance of two cell clusters is defined as the average decrease in the cell likelihood value if the cells from these two clusters are grouped as one cluster, as opposed to the original clustering. The lineage progression graph is built by adding transition edges between pairs of clusters in increasing magnitude of cluster distance until all clusters are connected to at least one other cluster, or based on user-specified criteria.

#### Single-cell transcriptional uncertainty

The last step in our analysis is to compute the final single-cell likelihood. Briefly, for each cell, we consider all edges in the lineage progression graph that are adjacent to the cell’s respective cluster, i.e. edges that eminate from or pointing to the cluster to which the cell belongs. The likelihood of a cell along an edge is evaluated by interpolating the likelihood values of the cell’s gene expression using the mRNA distributions from the two adjacent clusters. Each cell is then assigned to the edge along which its interpolated likelihood value is maximum, and the final cell likelihood is set to this maximum value. As mentioned above, the single-cell transcriptional uncertainty is evaluated as the negative logarithm of the cell likelihood value (NLL).

#### Pseudotimes calculation

For we can evaluate the pseudotimes for the cells according to the following procedure. First, a pseudotime is given to each cluster with a value between 0 (initial cell state) and 1 (final cell fate). Subsequently, we determine the linear fractional position of each cell along its respective edge at which its interpolated likelihood value is maximum (see *Single-cell transcriptional uncertainty*). The pseudotime of a cell is computed by a linear interpolation of the pseudotimes of the two clusters adjacent to its assigned edge according to the cell’s linear fractional position on this edge.

#### Epigenetic landscape reconstruction

To visualize the 3D transcriptional uncertainty landscape, we apply dimensional reduction techniques such as Principal Component Analysis (PCA) or t-SNE on the z-scored expression data, to project the gene expression of each individual cell on two dimensional axis, which gives the x-y axis of the landscape plot. For the z axis, we plot the NLL values. The transcriptional uncertainty landscape surface is reconstructed by estimating local approximation of individual cell 3D coordinates on a regular 30×30 grid by using a publicly available Matlab surface fitting code called GRIDFIT ^67^.

### Pre-processing and analysis of single-cell expression datasets

#### Bargaje et al. scRT-qPCR dataset

The dataset includes the expression profiles of 96 genes from 1896 single cells at 8 different time points (day 0, 1, 1.5, 2, 2.5, 3, 4, 5) during the differentiation of human pluripotent stem cells (iPSCs) into either mesodermal (M) or endodermal (En) fate ^35^. By employing CALISTA, we obtained five cell clusters and detected a bifurcation event, which gives rise to the two final cell fates. After lineage inference, we pseudotemporally ordered cells along the inferred differentiation paths (for more details, see ^40^).

#### Treutlein et al. scRNA-sequencing dataset

The dataset includes the gene expression profiles of 405 cells during reprogramming of mouse embryonic fibroblast (MEF) into a desired induced neural (iN) and an alternative myogenic (M) cell fates ^38^. We pre-processed the data using CALISTA to select the 40 most variable genes (10% of the number of cells) for the transcriptional uncertainty analysis. CALISTA identified four different subpopulations and successfully recovered the bifurcation event (for more details, see ^40^)

#### Richard et al. scRT-qPCR dataset

The dataset contains the expression profile of 91 genes measured from 389 cells at 6 distinct time points (0, 8, 24, 33, 48, 72 h) during the differentiation of primary chicken erythrocytic progenitor cells (T2EC) ^8^. Following CALISTA pre-processing step, we removed cells in which less than 75% of the genes are expressed. Then, we selected the subset of genes with at least one non-zero expression values. A total of 354 cells and 88 genes were considered in the transcriptional uncertainty analysis. Based on eigengap heuristics ^40,47^, we grouped cells into 6 optimal clusters and ordered cells along the inferred linear trajectory (see Supplementary Figure S7).

#### Stumpf et al. scRT-qPCR dataset

The dataset comprises the single-cell expression of 97 genes at 7 time points (0, 24, 48, 72, 96, 120, 168 h) during neural differentiation of mouse embryonic stem cells (E14 cell line) ^10^. In the data pre-processing, we excluded cells in which less than 70% of genes are expressed. Then, we selected genes with at least one non-zero expression values. A total of 276 cells and 93 genes were considered for for the transcriptional uncertainty analysis. Based on eigengap heuristics ^40^, we grouped cells into five optimal clusters and ordered cells along the inferred linear trajectory (Supplementary Figure S8).

#### Moussy et al. scRT-qPCR dataset

The single-cell expression dataset includes normalized Ct values for 91 genes in 435 cells captured at 5 distinct time points (0, 24, 48, 72, 96 h) during human cord blood-derived CD34+ differentiation ^12^. We employed CALISTA to group cells into 7 clusters, reconstruct the developmental trajectory and calculate pseudotimes (Supplementary Figure S9).

#### Guo et al. scRT-qPCR dataset

The dataset comprises the single-cell expression values of 48 genes from 387 individual cells isolated at 4 distinct developmental cell stages, from 8-cell stage mouse embryos to 64-blastocyst ^36^. By applying CALISTA, we identified seven different subpopulations along the differentiation process, and the inferred lineage hierarchy pinpointed two bifurcations events at 32-and 64-cell stage (Supplementary Figure S10). The timing of the lineage bifurcations coincides with two well-known branching points: one at 32-cell stage when totipotent cells differentiate into trophectoderm (TE) and inner cell mass (ICM), and another at 64-cell stage when ICM cells differentiate into primitive endoderm (PE) and epiblast (E).

#### Nestorowa et al. scRNA-sequencing dataset

The dataset comprises single-cell gene expression of 1656 cells from mouse hematopoietic stem cell differentiation ^39^. We pre-processed the data by removing genes with non-zero values in less than 10% of the cells. Then, we selected 433 most variable genes, which is 10% of the number of genes after the previous pre-processing step, for the transcriptional uncertainty analysis ^40^. We set the optimal number of clusters based on the original study ^39^, which reported six different subpopulations and two bifurcation events: the first one producing common myeloid progenitor (CMP) from lymphoid-primed multipotent progenitors (LMPP), and the second one generating granulocyte–monocyte progenitors (GMP) from megakaryocyte-erythroid progenitors (MEP) (Supplementary Figure S11).

#### Moignard et al. scRT-qPCR dataset

The dataset contains the single-cell expression level of 18 transcription factors measured in a total of 597 mouse bone marrow cells during hematopoietic differentiation. By applying CALISTA, we successfully identified the five subpopulations and the two branching points detected in the original study ^37^: long-term hematopoietic stem cells (HSC) differentiating into megakaryocyte–erythroid progenitors (PreM) or lymphoid-primed multipotent progenitors (LMPP); LMPP cells differentiating into granulocyte–monocyte progenitors (GMP) and common lymphoid progenitors (CLP) (for details see ^40^).

### Pairwise correlation analysis of transcriptional uncertainty and transcriptional burst size and frequency

We defined gene transcriptional burst size and burst frequency using the two-state model parameters, as follows:

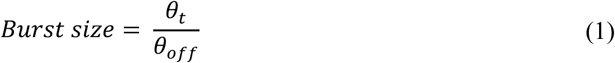

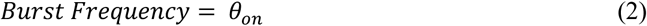

The burst sizes and burst frequencies are evaluated using the parameters 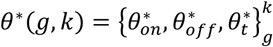 obtained from single-cell clustering analysis of CALISTA. Meanwhile, the average gene-wise NLL values for each single-cell cluster was computed as:

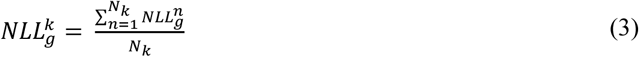

where 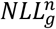 is the negative log-likelihood of cell *n* based only on the expression of gene *g*, and *N*_*k*_ is the total number of cells in cluster *k*.

## RNA VELOCITY ANALYSIS

Cells and genes were first filtered based on the pre-processing strategy in the original publication by La Manno and colleagues ^42^, which resulted in a total of 1720 cells and 1448 genes from human glutamatergic neurogenesis, and a total of 18140 cells and 2141 genes from mouse hippocampus dataset. We further reduced the number of genes to only the top 500 highly variable genes for the transcriptional uncertainty analysis. The cell cluster assignments generated by Velocyto – the algorithm for computing RNA velocity from the original publication – were considered, instead of using CALISTA. Based on the clustering, we employed CALISTA to generate the lineage progression and cell pseudotimes (Supplementary Figure S12). The RNA velocity and transcriptional uncertainty values for the top 500 genes were calculated by employing Velocyto and CALISTA, respectively. The cell-wise RNA velocity was set to the Euclidean norm of the vector of RNA velocities for each cell, while the cell-wise NLLs was computed according to:

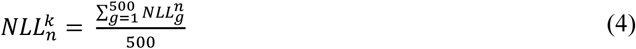

## CODE AVAILABILITY

A MATLAB and R version of CALISTA used in this study is freely available from the following website: https://www.cabselab.com/calista. All additional data are available from the corresponding author upon request.

## DATA AVAILABILITY

All the public single cell data sets analysed in this study are available from the original publications.

## ACKNOWLEDGEMENTS

We would like to thank all members of the SBDM team for lively discussions. We also thank the BioSyL Federation and the LabEx Ecofect (ANR-11-LABX-0048) of the University of Lyon for inspiring scientific events.

