## Supplementary Figure for "Universality of cell differentiation trajectories revealed by a reconstruction of transcriptional uncertainty landscapes from single-cell transcriptomic data"

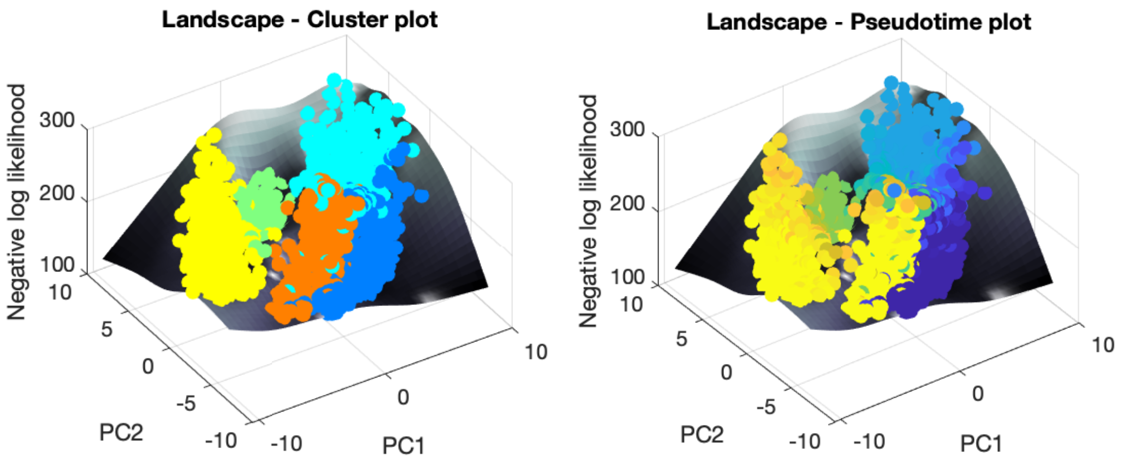


**Figure S1. Cell likelihood landscape plots for Bargaje et al. data [1] considering empirical gene distributions.** Single cells were pseudotemporally ordered along the inferred differentiation trajectories based on the maximum likelihood optimization by applying linear interpolation over the empirical (observed) distributions. Landscape surfaces were depicted for cell clusters (*left*) and pseudotimes from blue to yellow (*right*).

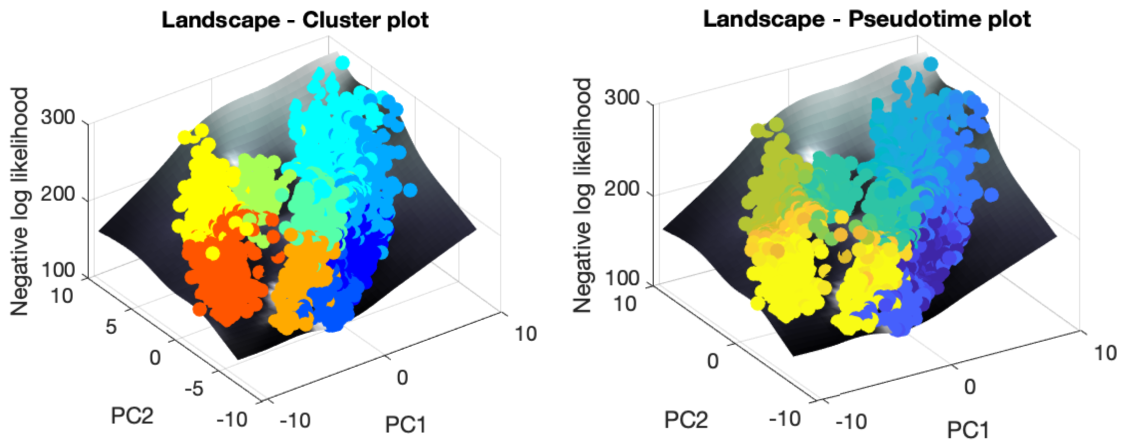


**Figure S2. Cell likelihood landscape plots for Bargaje et al. data [1] considering nine cell clusters.** The number of clusters were set to nine based on eigengap plot [2]. Expression data were analyzed by using CALISTA for cell clustering, lineage inference, cell ordering and landscape plotting. Landscape surfaces were depicted for cell clusters (*left*) and pseudotimes from blue to yellow (*right*).

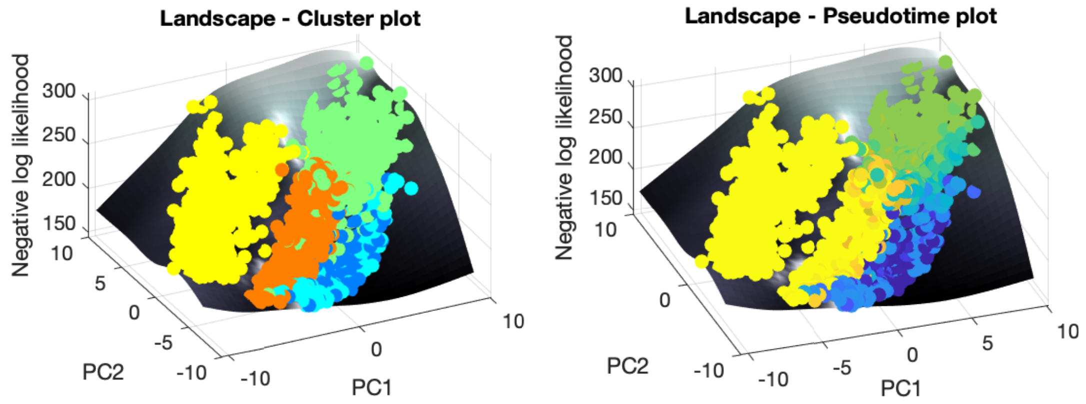


**Figure S3. Cell likelihood landscape plots for Bargaje et al. data [1] considering SIMLR cell clusters.** Subpopulations were identified by applying SIMLR [3] cell clustering. Epigenetic landscapes were reconstructed after inferring the lineage relationships and pseudotemporal ordering by CALISTA. Landscape surfaces were depicted for cell clusters (*left*) and pseudotimes from blue to yellow (*right*).

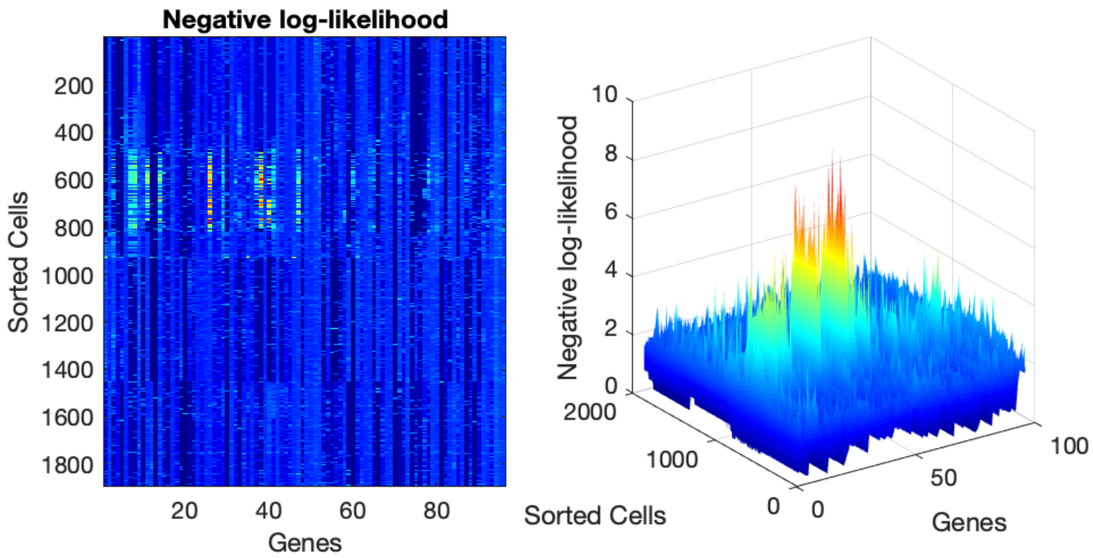


**Figure S4. Contribution of genes to single-cell transcriptional uncertainty in Bargaje et al. data [1].** Heatmap (*left*) and 3D plot (*right*) of NLL value of each gene in each cell (for further details see Methods in [2]). Cells were sorted in increasing order of cluster numbers (e.g. cells in cluster 1, cells in cluster 2, etc.). Peaks in NLL values can be observed in specific gene sets.

| Index | Name | P-value | Adjusted p-value | Z-score | Combined score |
|---|---|---|---|---|---|
| 1 | Cardiac Progenitor Differentiation_Homo sapiens_WP2406 | 1.771e-17 | 1.204e-15 | -1.93 | 74.44 |
| 2 | Endoderm Differentiation_Homo sapiens_WP2853 | 5.847e-14 | 1.988e-12 | -1.94 | 59.11 |
| 3 | Mesodermal Commitment Pathway_Homo sapiens_WP2857 | 1.913e-11 | 4.335e-10 | -1.79 | 44.21 |
| 4 | Heart Development_Homo sapiens_WP1591 | 2.466e-10 | 3.354e-9 | -1.77 | 39.17 |
| 5 | Heart Development_Mus musculus_WP2067 | 1.748e-10 | 2.972e-9 | -1.71 | 38.48 |
| 6 | PluriNetWork_Mus musculus_WP1763 | 0.000002142 | 0.00002427 | -2.06 | 26.87 |
| 7 | Adipogenesis genes_Mus musculus_WP447 | 0.000002745 | 0.00002482 | -1.74 | 22.24 |
| 8 | Adipogenesis_Homo sapiens_WP236 | 0.000002920 | 0.00002482 | -1.71 | 21.79 |
| 9 | Differentiation Pathway_Homo sapiens_WP2848 | 0.000007108 | 0.00005370 | -1.77 | 21.01 |
| 10 | MicroRNAs in Cardiomyocyte Hypertrophy_Mus musculus_WP1560 | 0.00003201 | 0.0002177 | -1.80 | 18.62 |

**Figure S5. Pathway enrichment analysis for Bargaje et al. data [1].** The analysis was performed by EnrichR [4] on the gene set identified for cluster 2 (see main text and Supplementary Table S1).

| Index | Name | P-value | Adjusted p-value | Z-score | Combined score |
|---|---|---|---|---|---|
| 1 | Mesodermal Commitment Pathway_Homo sapiens_WP2857 | 1.332e-8 | 3.330e-7 | -1.85 | 33.50 |
| 2 | Endoderm Differentiation_Homo sapiens_WP2853 | 0.000002370 | 0.00002963 | -1.94 | 25.13 |
| 3 | Heart Development_Homo sapiens_WP1591 | 0.0001500 | 0.0009374 | -1.81 | 15.92 |
| 4 | Heart Development_Mus musculus_WP2067 | 0.0001313 | 0.0009374 | -1.75 | 15.66 |
| 5 | Cardiac Progenitor Differentiation_Homo sapiens_WP2406 | 0.0001910 | 0.0009549 | -1.79 | 15.33 |
| 6 | Neural Crest Differentiation_Homo sapiens_WP2064 | 0.0006932 | 0.002888 | -1.79 | 13.03 |
| 7 | Adipogenesis genes_Mus musculus_WP447 | 0.001110 | 0.003576 | -1.74 | 11.81 |
| 8 | Adipogenesis_Homo sapiens_WP236 | 0.001144 | 0.003576 | -1.71 | 11.58 |
| 9 | Differentiation of white and brown adipocyte _Homo sapiens_WP2895 | 0.009958 | 0.02766 | -1.61 | 7.40 |
| 10 | Striated Muscle Contraction_Mus musculus_WP216 | 0.01629 | 0.03475 | -1.63 | 6.71 |

**Figure S6. Pathway enrichment analysis for Bargaje et al. data [1].** The analysis was performed by EnrichR [4] on the gene set identified for cluster 3 (see main text and Supplementary Table S1).

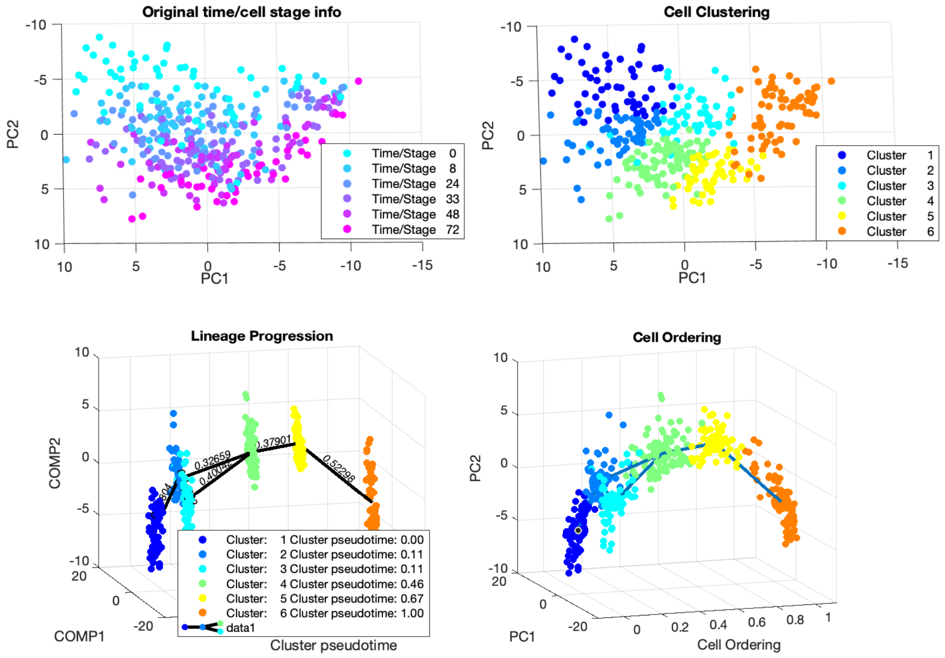


**Figure S7. CALISTA analysis of Richard et al. data [5].** The plots show the expression data based on the original time info (*top left*), cell clustering assignments (*top right*), inferred lineage relationships (*bottom left*), and cell ordering (*bottom right*).

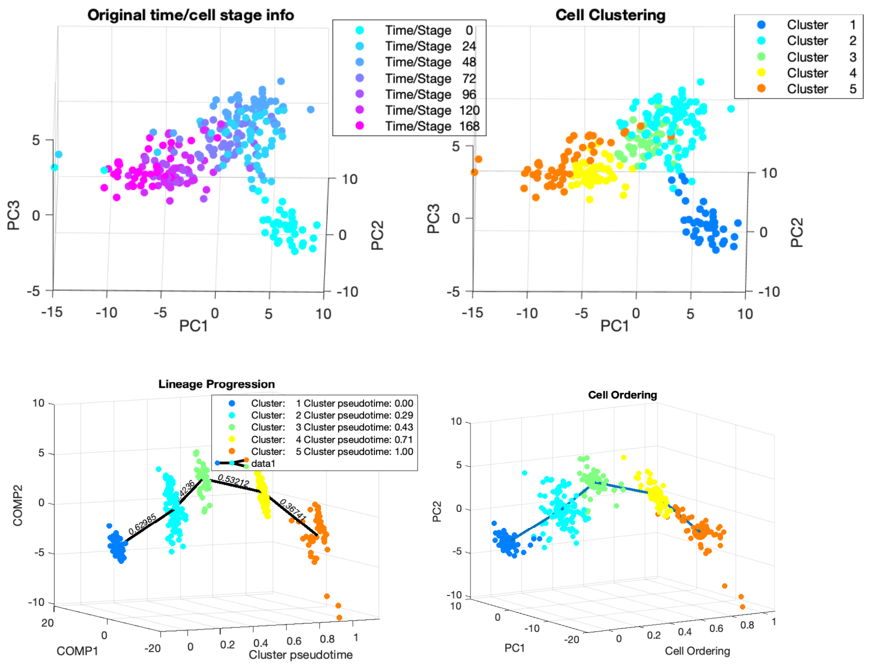


**Figure S8. CALISTA analysis of Stumpf et al. data [6].** The plots show the expression data based on the original time info (*top left*), cell clustering assignments (*top right*), inferred lineage relationships (*bottom left*), and cell ordering (*bottom right*).

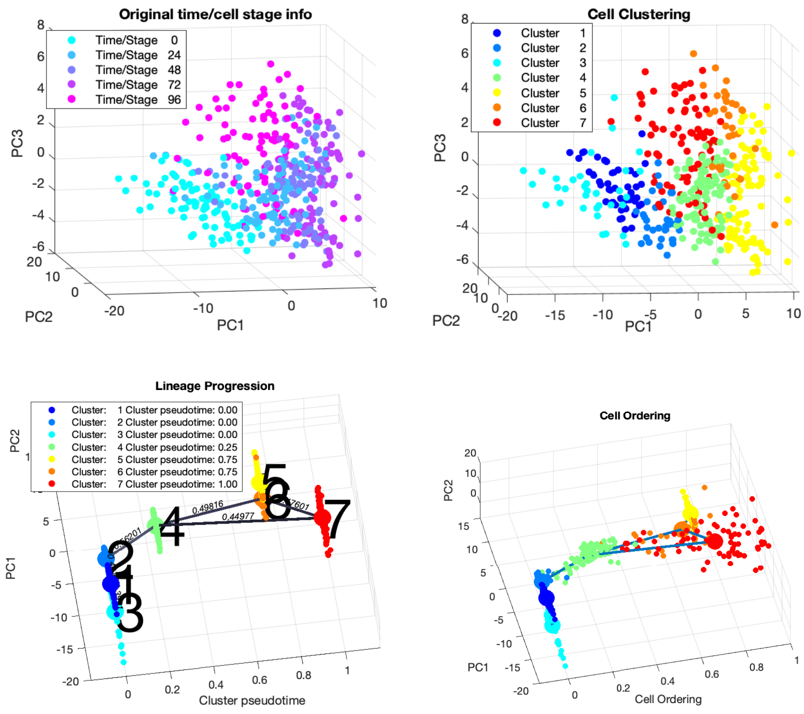


**Figure S9. CALISTA analysis of Moussy et al. data [7].** The plots show the expression data based on the original time info (*top left*), cell clustering assignments (*top right*), inferred lineage relationships (*bottom left*), and cell ordering (*bottom right*). An edge connecting cluster 2 and 7 was manually removed because it was inconsistent with the cell capture time information.

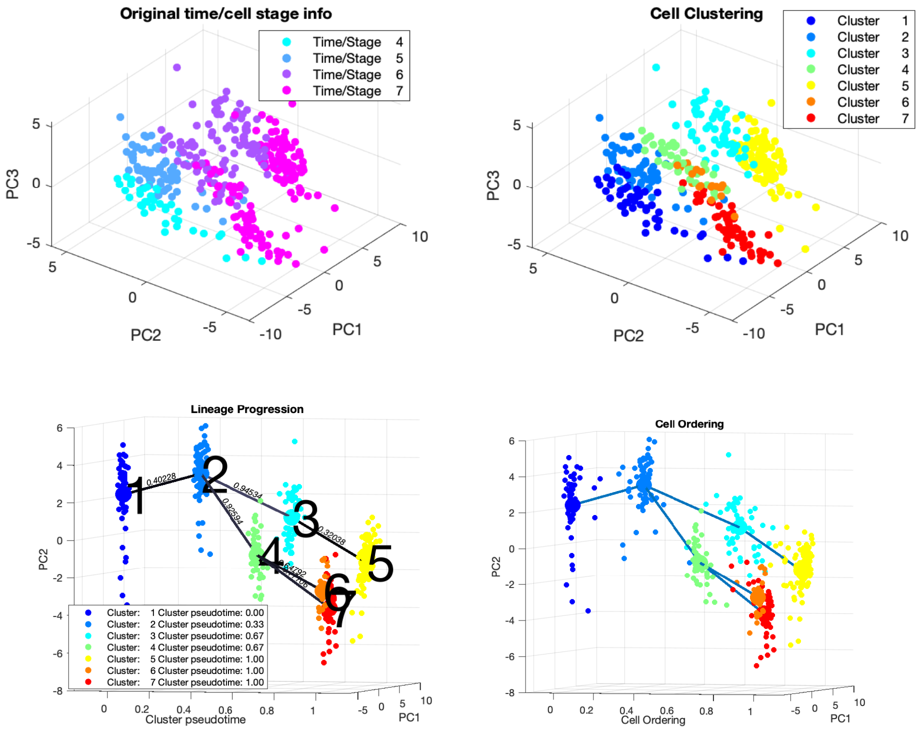


**Figure S10. CALISTA analysis of Guo et al. data [8].** The plots show the expression data based on the original cell stage info (*top left*), cell clustering assignments (*top right*), inferred lineage relationships (*bottom left*), and cell ordering (*bottom right*). An edge connecting cluster 1 and 4 was manually removed as the edge was inconsistent with the cell stage information.

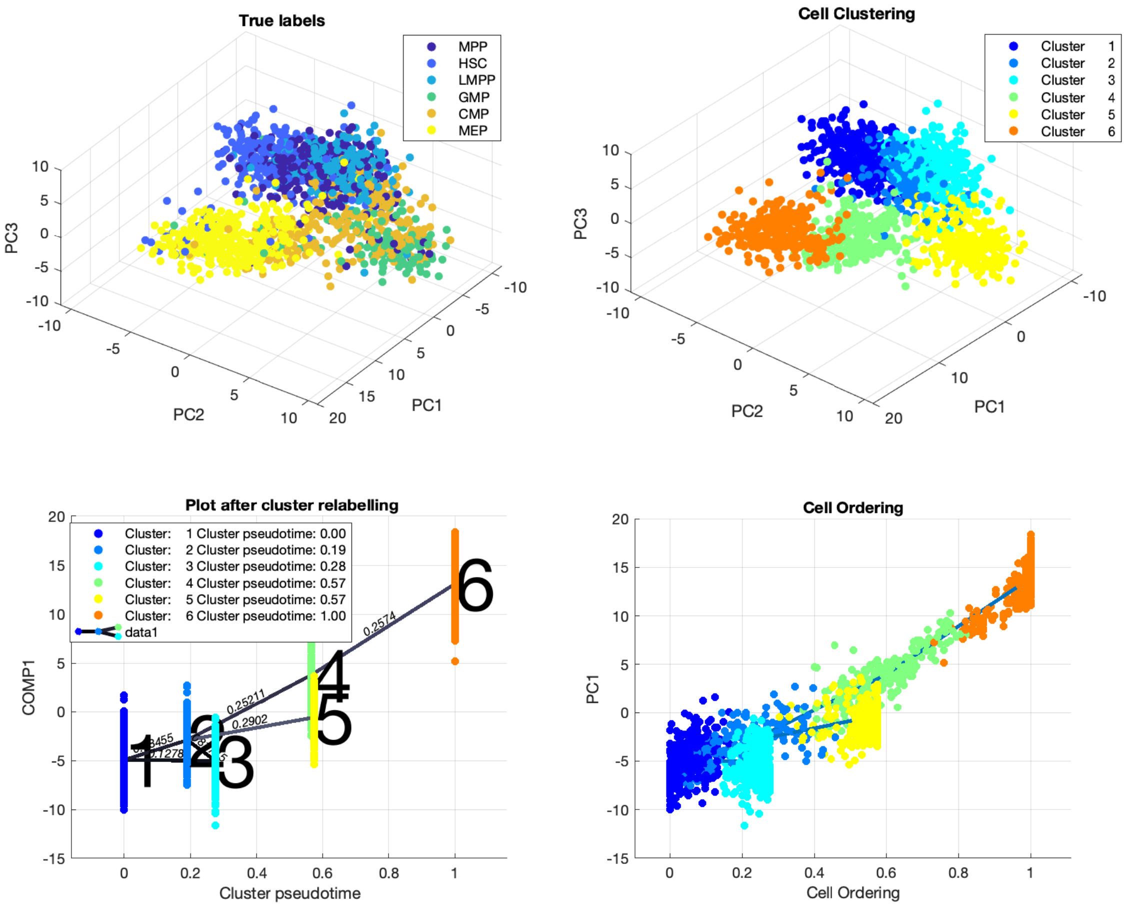


**Figure S11. CALISTA analysis of Nestorowa et al. data [9].** The plots show the expression data based on the original cell labels (*top left*), cell clustering assignments (*top right*), inferred lineage relationships (*bottom left*), and cell ordering (*bottom right*).

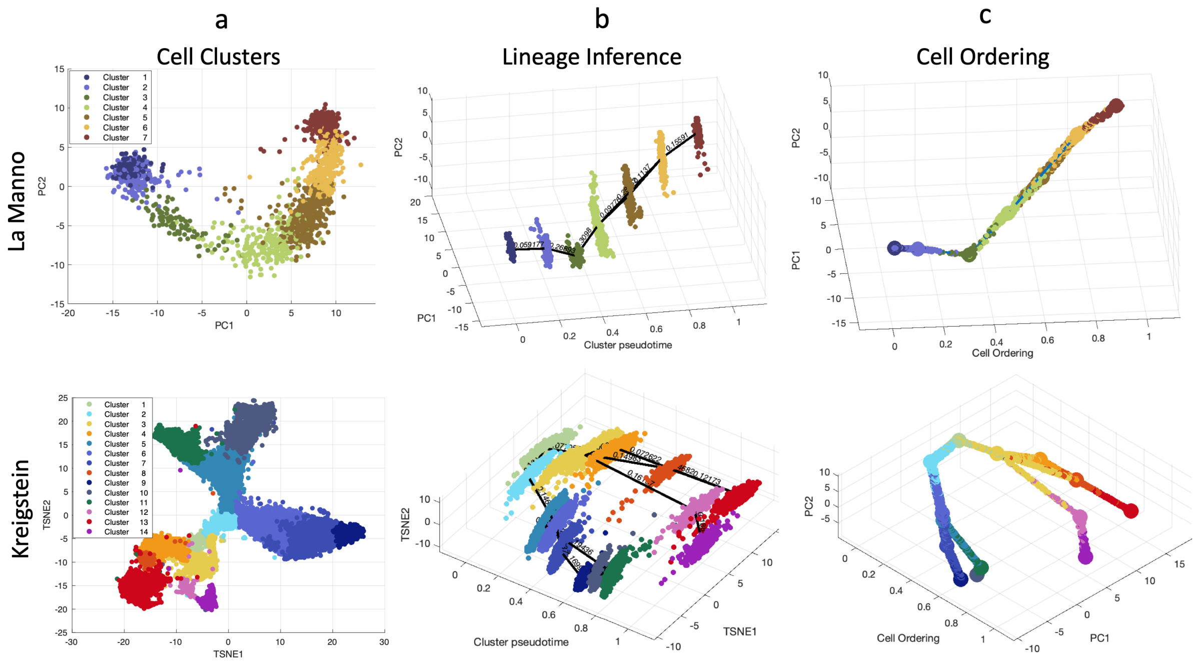


**Figure S12 Single-cell expression data analysis.** (Top row) Human glutamatergic neurogenesis in La Manno et al. study [10]. (Bottom row) Mouse hippocampal neurogenesis in Kreigstein et al. study [11]. (First column) Cell clustering assignments evaluated from Velocyto. Lineage progresison (Second column) and cell ordering (Third column) were predicted by CALISTA.  PC: principal component. TSNE: t-Distributed Stochastic Neighbor Embedding component. The colors in the first column indicate the cell clusters.

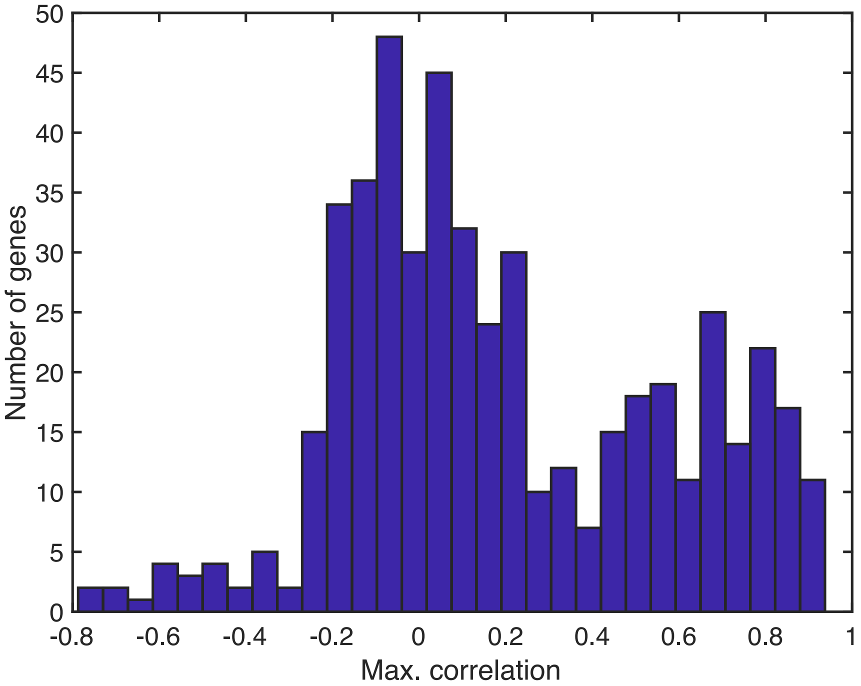


**Figure S13 Gene-wise cross correlation analysis between RNA velocity and single-cell transcriptional analysis (glutamatergic neurogenesis)[10].** Histogram of the maximum correlation between RNA velocity and time-shifted (advanced) single-cell transcriptional uncertainty for each of the top 500 most variable genes. First, for each gene, the RNA velocity and transcriptional uncertainty of the cells were first sorted in increasing cell pseudotime. Then, we performed cross correlation to determine the time difference that gives the maximum correlation between the RNA velocity and the delayed transcriptional uncertainty. The 25$^{th}$, 50$^{th}$ and 75$^{th}$ percentile of the correlations are −0.075, 0.112, and 0.510.
