## Supplementary figures and images for "Universality of cell differentiation trajectories revealed by a reconstruction of transcriptional uncertainty landscapes from single-cell transcriptomic data"

### Supplementary File S1

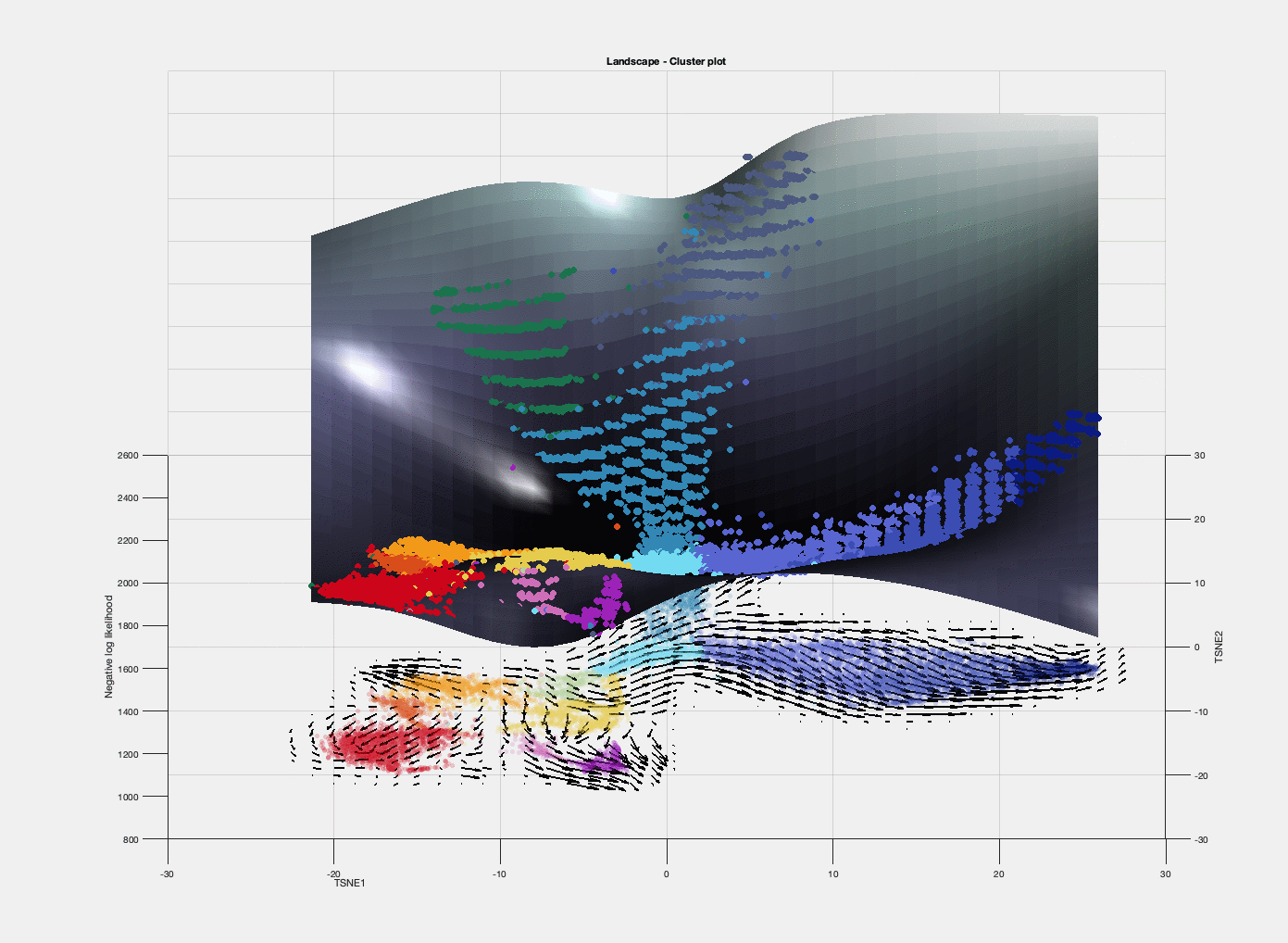

### Supplementary File S2

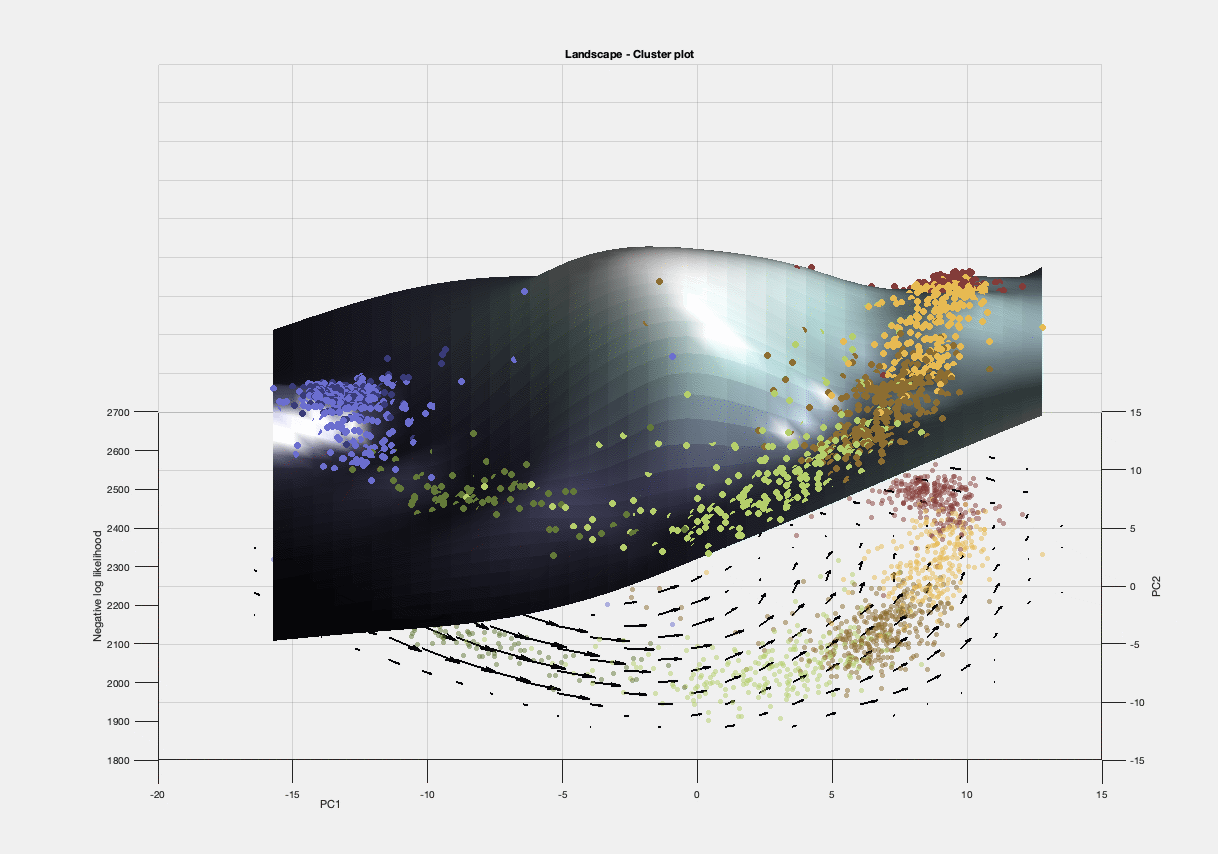
